# ESCRT recruitment to mRNA-encoded SARS-CoV-2 spike induces virus-like particles and enhanced antibody responses

**DOI:** 10.1101/2022.12.26.521940

**Authors:** Magnus A. G. Hoffmann, Zhi Yang, Kathryn E. Huey-Tubman, Alexander A. Cohen, Priyanthi N. P. Gnanapragasam, Leesa M. Nakatomi, Kaya N. Storm, Woohyun J. Moon, Paulo J.C. Lin, Pamela J. Bjorkman

**Affiliations:** Division of Biology and Biological Engineering, California Institute of Technology, Pasadena, CA 91125, USA; Acuitas Therapeutics, Vancouver, BC, V6T 1Z3, CANADA

## Abstract

Prime-boost regimens for COVID-19 vaccines elicit poor antibody responses against Omicron-based variants and employ frequent boosters to maintain antibody levels. We present a natural infection-mimicking technology that combines features of mRNA- and protein nanoparticle-based vaccines through encoding self-assembling enveloped virus-like particles (eVLPs). eVLP assembly is achieved by inserting an ESCRT- and ALIX-binding region (EABR) into the SARS-CoV-2 spike cytoplasmic tail, which recruits ESCRT proteins to induce eVLP budding from cells. Purified spike-EABR eVLPs presented densely-arrayed spikes and elicited potent antibody responses in mice. Two immunizations with mRNA-LNP encoding spike-EABR elicited potent CD8+ T-cell responses and superior neutralizing antibody responses against original and variant SARS-CoV-2 compared to conventional spike-encoding mRNA-LNP and purified spike-EABR eVLPs, improving neutralizing titers >10-fold against Omicron-based variants for three months post-boost. Thus, EABR technology enhances potency and breadth of vaccine-induced responses through antigen presentation on cell surfaces and eVLPs, enabling longer-lasting protection against SARS-CoV-2 and other viruses.

## Introduction

mRNA vaccines emerged during the COVID-19 pandemic as an ideal platform for the rapid development of effective vaccines (Corbett et al., 2020). Currently approved SARS-CoV-2 mRNA vaccines encode the viral spike (S) trimer (Zheng et al., 2022), the primary target of neutralizing antibodies during natural infections (Chen et al., 2022). Clinical studies have demonstrated that mRNA vaccines are highly effective, preventing >90% of symptomatic and severe SARS-CoV-2 infections (Baden et al., 2021; Polack et al., 2020) through both B and T cell responses (Kent et al., 2022). mRNA vaccines in part mimic an infected cell since expression of S within cells that take up S-encoding mRNAs formulated in lipid nanoparticles (LNP) (Hogan and Pardi, 2022) results in cell surface expression of S protein to stimulate B cell activation. Translation of S protein inside the cell also provides viral peptides for presentation on MHC class I molecules to cytotoxic T cells, which does not commonly occur in protein nanoparticle-based vaccines (Rock et al., 2016) that resemble the virus by presenting dense arrays of S protein; e.g., the Novavax NVX-CoV2373 vaccine (Heath et al., 2021; Keech et al., 2020). However, comparisons to COVID-19 mRNA vaccines showed that NVX-CoV2373 elicits comparable neutralizing antibody titers (Karbiener et al., 2022; Zhang et al., 2022), the main immune correlate of vaccine-induced protection (Barouch, 2022), suggesting that potent B cell activation can be achieved through presentation of viral surface antigens on cell surfaces or virus-resembling nanoparticles. Achieving higher antibody neutralization titers is desirable as antibody levels contract substantially over a period of several months (Zhang et al., 2022), and SARS-CoV-2 variants of concern (VOCs) that are less sensitive to antibodies elicited by vaccines or natural infection have been emerging (Chen et al., 2021; Hachmann et al., 2022; Wu et al., 2021). An optimal vaccine might therefore combine attributes of both mRNA- and protein nanoparticle-based vaccines by delivering a genetically encoded S protein that gets presented on cell surfaces and induces self-assembly and release of S-presenting nanoparticles.

Here, we describe a novel technology that engineers membrane proteins to induce self-assembly of enveloped virus-like particles (eVLPs) that bud from the cell surface. This is accomplished for the SARS-CoV-2 S protein by inserting a short amino acid sequence (termed an ESCRT- and ALIX-binding region or EABR) (Lee et al., 2008) at the C-terminus of its cytoplasmic tail to recruit host proteins from the <u>e</u>ndosomal <u>s</u>orting <u>c</u>omplex <u>r</u>equired for transport (ESCRT) pathway. Many enveloped viruses recruit ESCRT-associated proteins such as TSG101 and/or ALIX through capsid or other interior viral structural proteins during the budding process (McCullough et al., 2018; Votteler and Sundquist, 2013). Thus, fusing the EABR to the cytoplasmic tail of a viral glycoprotein or other membrane protein directly recruits TSG101 and ALIX, bypassing the need for co-expression of other viral proteins for eVLP self-assembly. Cryo-electron tomography (cryo-ET) showed dense coating of spikes on purified S-EABR eVLPs, and direct injections of the eVLPs elicited potent neutralizing antibody responses in mice. Finally, we demonstrate that an mRNA vaccine encoding the S-EABR construct elicited at least 5-fold higher neutralizing antibody responses against SARS-CoV-2 and VOCs in mice than a conventional S-encoding mRNA vaccine or purified S-EABR eVLPs. These results demonstrate that mRNA-mediated delivery of S-EABR eVLPs elicits superior antibody responses, suggesting that dual presentation of viral surface antigens on cell surfaces and on extracellular eVLPs has the potential to enhance the effectiveness of COVID-19 mRNA vaccines.

## Results

### ESCRT recruitment to the spike cytoplasmic tail induces eVLP assembly

To evaluate the hypothesis that direct recruitment of ESCRT proteins to the cytoplasmic tail of a SARS-CoV-2 S protein could result in self-assembly and budding of eVLPs, we fused EABRs derived from different sources to the truncated cytoplasmic tail of the S protein, separated from its C-terminus by a short Gly-Ser linker (Figures 1A and 1B). The S protein contained the D614G substitution (Korber et al., 2020), a furin cleavage site, two proline substitutions (2P) in the S2 subunit to stabilize the prefusion conformation (Pallesen et al., 2017), and the C-terminal 21 residues were truncated to optimize cell surface expression by removing an endoplasmic reticulum (ER)-retention signal (ΔCT) (McBride et al., 2007) (Figure 1B). We evaluated the EABR fragment from the human CEP55 protein that binds TSG101 and ALIX during cytokinesis (Lee et al., 2008) (Figure 1B). For comparisons, viral late domains that recruit early ESCRT proteins during the viral budding process were obtained from the Equine Infectious Anemia Virus (EIAV) p9 protein (Fisher et al., 2007), residues 1-44 of the Ebola virus (EBOV) VP40 protein (Madara et al., 2015), and the HIV-1 p6 protein (Fujii et al., 2009) (Figure S1A). We hypothesized that eVLP production could be enhanced by preventing endocytosis of EABR-fusion proteins to extend the duration that proteins remain at the plasma membrane to interact with ESCRT proteins. We therefore added an endocytosis prevention motif (EPM), a 47-residue insertion derived from the murine Fc gamma receptor FcgRBII-B1 cytoplasmic tail (Figures 1A and 1B) that tethers FcgRBII-B1 to the cytoskeleton to prevent coated pit localization and endocytosis (Miettinen et al., 1989).

**Figure 1.**
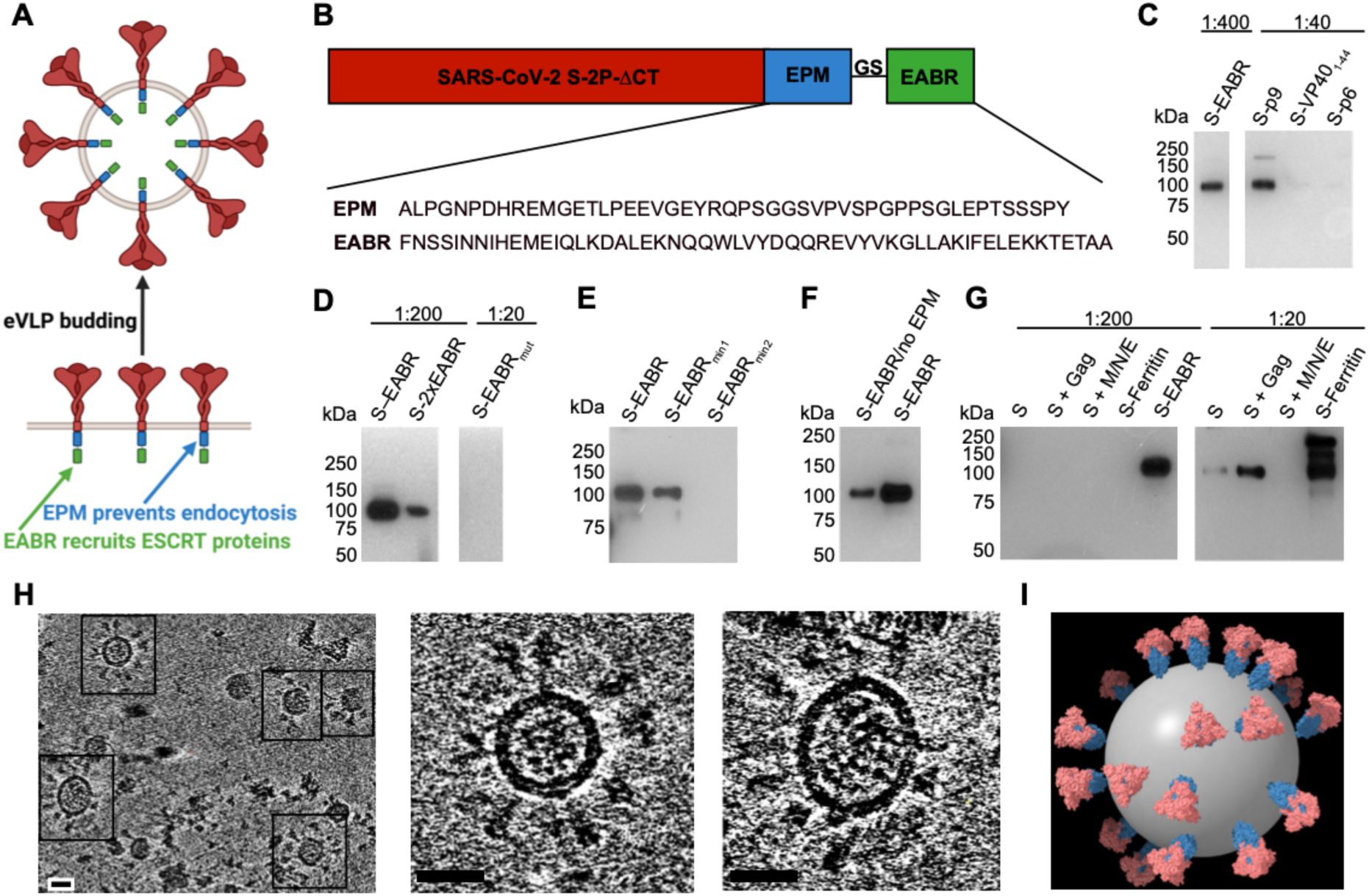
EABR insertion into the cytoplasmic tail of membrane proteins results in eVLP budding and release. (A) Schematic of membrane-bound SARS-CoV-2 S proteins on the cell surface containing cytoplasmic tail EPM and EABR insertions that induce budding of an eVLP comprising a lipid bilayer with embedded S proteins. (B) Sequence information for S-EABR construct. Top: The SARS-CoV-2 S protein (including a furin cleavage site, 2P stabilizing substitutions, the D614G substitution, and ΔCT, a cytoplasmic tail deletion) is fused to an EPM sequence, a (Gly)_3_Ser (GS) spacer, and an EABR sequence. EPM = Endocytosis prevention motif. GS = (Gly)_3_Ser linker. EABR = ESCRT- and ALIX-binding region. Bottom: EPM and EABR sequence information. (C-G) Western blot analysis detecting SARS-CoV-2 S1 protein on eVLPs purified by ultracentrifugation on a 20% sucrose cushion from transfected Expi293F cell culture supernatants. (C) Cells were transfected with S-EABR, S-p9, S-VP40_1-44_, or S-p6 constructs. The purified S-EABR eVLP sample was diluted 1:400 (left), while S-p9, S-VP40_1-44_, and S-p6 samples were diluted 1:40 (right). Comparison of band intensities between lanes suggest that the S-EABR eVLP sample contained ∼10-fold higher levels of S1 protein than the S-p9 sample and >10-fold higher levels than the S-VP40_1-44_ and S-p6 samples. (D) Cells were transfected with S-EABR, S-2xEABR (left) or S-EABR_mut_ constructs (right). Purified S-EABR and S-2xEABR eVLP samples were diluted 1:200, while the S-EABR_mut_ sample was diluted 1:20. (E) Cells were transfected with S-EABR, S-EABR_min1_, or S-EABR_min2_ constructs. Purified eVLP samples were diluted 1:200. (F) Cells were transfected with S-EABR/no EPM or S-EABR constructs. Purified eVLP samples were diluted 1:200. (G) Cells were transfected to express S alone, S plus the HIV-1 Gag protein, S plus the SARS-CoV-2 M, N, and E proteins, an S-ferritin fusion protein, or S-EABR. Purified eVLP samples were diluted 1:200 (left) or 1:20 (right). Comparison of band intensities between lanes suggest that the S-EABR eVLP sample contained >10-fold higher levels of S1 protein than S alone, S plus Gag, and S plus M, N, E. (H) Computationally-derived tomographic slices (8.1 nm) of S-EABR eVLPs derived from cryo-ET imaging of S-EABR eVLPs purified from transfected cell culture supernatants by ultracentrifugation on a 20% sucrose cushion and SEC. Left: Representative eVLPs are highlighted in boxes. Middle and right: Close-ups of individual eVLPs. Scale bars = 30 nm. (I) Model of a representative S-EABR eVLP derived from a cryo-ET reconstruction (Movie S1). Coordinates of an S trimer (PDB 6VXX) (Walls et al., 2020) were fit into protruding density on the best resolved half of an eVLP and the remainder of the eVLP was modeled assuming a similar distribution of trimers. The position of the lipid bilayer is shown as a 55 nm gray sphere.

The abilities of the S-EABR, S-p9, S-VP40_1-44_, and S-p6 constructs to generate eVLPs were evaluated by transfecting Expi293F cells and measuring eVLP production in supernatants from which eVLPs were purified by ultracentrifugation on a 20% sucrose cushion. Western blot analysis showed that the highest S protein levels were detected for the S-EABR construct, suggesting that the CEP55 EABR induced efficient self-assembly of S-containing eVLPs (Figures 1C and S1B). At a sample dilution of 1:400, the S-EABR construct produced a similarly intense band compared to the S-p9 construct at a 1:40 dilution, suggesting that S protein levels were ∼10-fold higher. The CEP55 EABR binds both ALIX and TSG101 (Lee et al., 2008), whereas EIAV p9 only binds ALIX (Fisher et al., 2007), suggesting that optimal recruitment of both ESCRT proteins is required for efficient eVLP assembly. The S-p6 and S-VP40_1-44_ samples contained little or no S protein suggesting that eVLP assembly was inefficient, possibly resulting from lower affinities for ESCRT proteins (Figures 1C and S1B).

We further characterized the S-EABR construct by experimenting with different EABR sequences (Figure S1A), finding that addition of a second EABR domain (S-2xEABR) reduced eVLP production (Figure 1D). To investigate whether S-EABR eVLP assembly is dependent on ESCRT recruitment, we generated S-EABR_mut_ by substituting an EABR residue (Tyr187 in CEP55) that is essential for interacting with ALIX (Lee et al., 2008) (Figure S1A). While the purified S-EABR eVLP sample produced an intense band at a 1:200 dilution, no band was detected for S-EABR_mut_ at a 1:20 dilution, suggesting that eVLP production was abrogated for S-EABR_mut_ and highlighting the importance of ALIX recruitment for eVLP assembly (Figure 1D). To identify the minimal EABR sequence required for eVLP assembly, we designed S constructs fused to the complete EABR domain (CEP55_170-213_), EABR_min1_ (CEP55_180-_

_213_), and EABR_min2_ (CEP55_180-204_) (Figure S1a). While S-EABR eVLP yields were diminished for EABR_min2_, production efficiency was retained for EABR_min1_ (Figure 1E). To assess the effects of the EPM within the cytoplasmic tail of the S-EABR construct, we evaluated eVLP production for an S-EABR construct that did not include the EPM. Western blot analysis demonstrated that increased amounts of S protein were detected after eVLP purification from cells transfected with S-EABR compared to S-EABR/no EPM, suggesting that the EPM enhances eVLP production (Figure 1F).

We also compared the S-EABR construct to other eVLP approaches (Martins et al., 2022) that require co-expression of S protein with structural viral proteins, such as HIV-1 Gag (Hoffmann et al., 2020) or the SARS-CoV-2 M, N, and E proteins (Syed et al., 2021). Western blot analysis showed that purified S-EABR eVLP fractions contained at least 10-fold more S protein than eVLPs produced by co-expression of S and Gag or S, M, N, and E (Figure 1G), suggesting that S-EABR eVLPs assemble and/or incorporate S proteins more efficiently than the other eVLP approaches. Purified S-EABR eVLPs also contained higher levels of S protein compared to S-ferritin nanoparticles purified from transfected cell supernatants, which have been shown to elicit potent immune responses in animal models (Joyce et al., 2021; Powell et al., 2021) (Figure 1G).

3D reconstructions derived from cryo-ET showed purified S-EABR eVLPs with diameters ranging from 40 - 60 nm that are surrounded by a lipid bilayer and the majority of which were densely coated with spikes (Figures 1H and 1I; Movie S1). To estimate the number of S trimers, we counted trimer densities in ∼4 nm computational tomographic slices of individual eVLPs, finding ∼10-40 spikes per particle that were heterogeneously distributed on the surface of eVLPs. The upper limit of the number of spikes on eVLPs roughly corresponds to spike numbers on larger SARS-CoV-2 virions (>100 nm in diameter) (Ke et al., 2020); thus, the spike densities on the majority of eVLPs exceed those on authentic viruses. Spikes on eVLPs were separated by distances of ∼20-26 nm (measured between the centers of trimer apexes) for densely coated particles (Figures 1H and 1I). To assess the general applicability of the EABR approach, we also generated EABR eVLPs for HIV-1 Env, which produced eVLPs with higher Env content than co-expression of Env and HIV-1 Gag (Figure S1C), and for the multi-pass transmembrane protein CCR5 (Figure S1D). Taken together, these results are consistent with efficient incorporation of S proteins into S-EABR eVLPs that are released from transfected cells and suggest that the EABR technology can be applied to a wide range of membrane proteins.

### S-EABR eVLPs induce potent antibody responses in immunized mice

The potential of purified S-EABR eVLPs as a vaccine candidate against SARS-CoV-2 was evaluated in C57BL/6 mice (Figure 2A). S-EABR eVLPs were purified from transfected cell supernatants by ultracentrifugation on a 20% sucrose cushion followed by size exclusion chromatography (SEC), and S protein concentrations were determined by quantitative Western blot analysis (Figures S2A and S2B). For a 100 mL transfection of Expi293F cells, purified S-EABR eVLPs from supernatants contained ∼250-500 µg S protein. Immunizations with S-EABR eVLPs were compared to purified soluble S and to soluble S covalently attached to SpyCatcher-mi3 protein nanoparticles (S-mi3) (Keeble et al., 2019). 0.1 µg doses (calculated based on S protein content) were administered by subcutaneous injections on days 0 and 28 for all immunogens in the presence of Sigma adjuvant (Figure 2A), and we evaluated serum antibody responses by enzyme-linked immunosorbent assays (ELISAs) and in vitro pseudovirus neutralization assays. After the prime, S-EABR eVLPs elicited robust antibody binding and neutralization responses in all mice against SARS-CoV-2 (WA1 variant including the D614G substitution (WA1/D614G)), similar to titers elicited by S-mi3 (Figures 2B and 2C). In contrast, no neutralizing antibody responses were detected for soluble S protein immunization after the prime. Neutralizing antibody titers elicited by S-EABR eVLPs and S-mi3 increased by >10-fold after boosting and were >20-fold higher than titers measured for soluble S (Figure 2C). S-EABR eVLPs elicited potent antibody responses targeting the receptor-binding domain (RBD) of the S protein (Figure S2C), a primary target of anti-SARS-CoV-2 neutralizing antibodies (Kleanthous et al., 2021). Serum responses were also evaluated against authentic SARS-CoV-2 by plaque reduction neutralization tests (PRNTs), showing robust neutralizing activity against SARS-CoV-2 WA1 (Figure S2D). Neutralization titers dropped ∼4-fold and ∼2-fold against the SARS-CoV-2 Beta and Delta variants, respectively, consistent with studies of licensed vaccines that encode the SARS-CoV-2 WA1 S protein (van Gils et al., 2022). These results demonstrate that purified S-EABR eVLPs elicit potent immune responses in vivo and represent an alternative technology for producing nanoparticle-based vaccines that does not involve detergent-mediated cell lysis and separation of membrane protein antigens from cell lysates, as required for protein nanoparticle vaccines such as NVX-CoV2373, a COVID-19 vaccine (Heath et al., 2021; Keech et al., 2020), or FluBlok, an influenza vaccine (Cox and Hollister, 2009).

**Figure 2.**
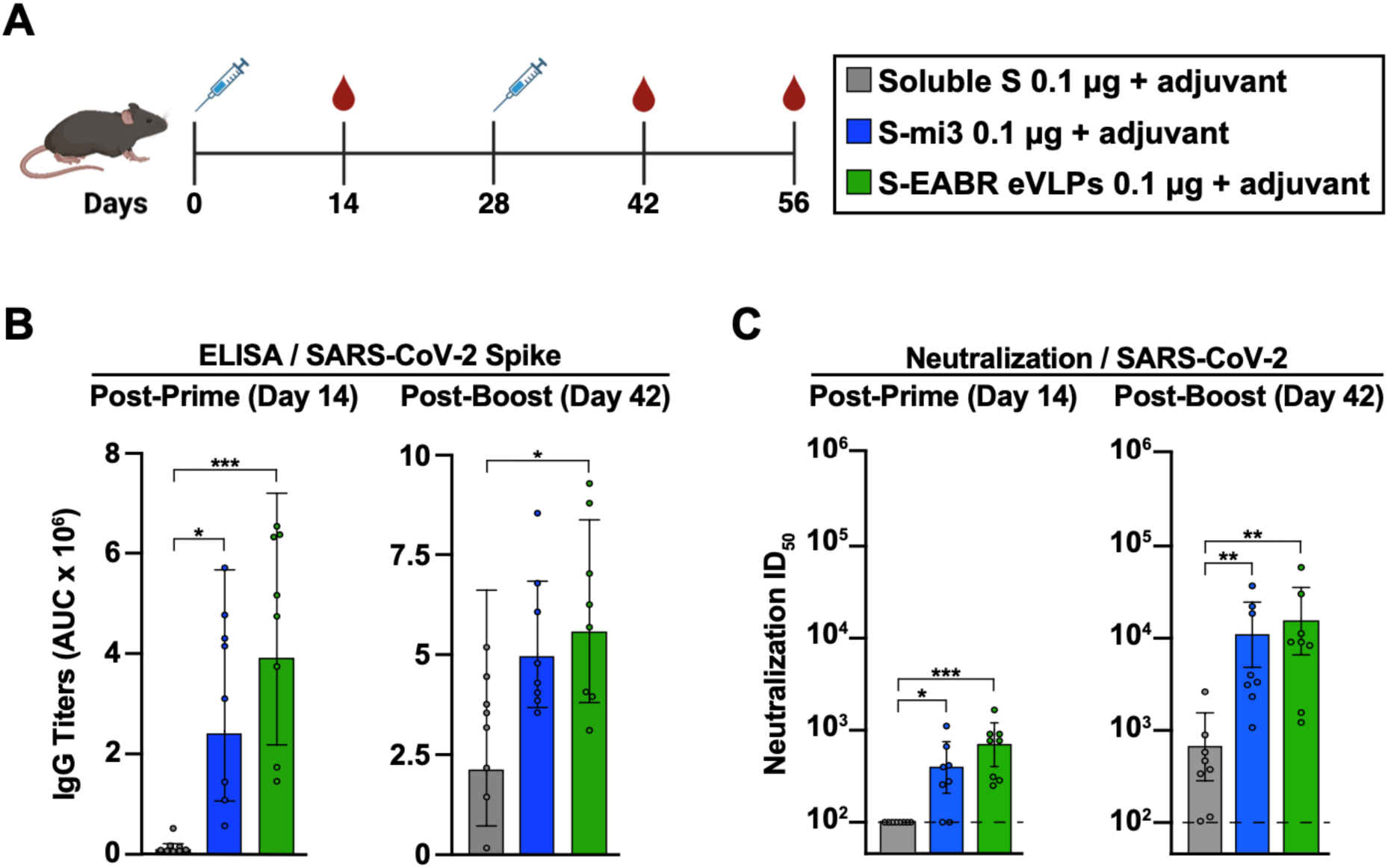
Purified S-EABR eVLPs induce potent antibody responses in mice. (A) Immunization schedule. C57BL/6 mice were immunized with soluble S (purified S trimer) (gray), S-mi3 (S trimer ectodomains covalently attached to mi3, a 60-mer protein nanoparticle) (blue), or S-EABR eVLPs (green). (B-C) ELISA and neutralization data from the indicated time points for antisera from individual mice (colored circles) presented as the geometric mean (bars) and standard deviation (horizontal lines). ELISA results are shown as area under the curve (AUC); neutralization results are shown as half-maximal inhibitory dilutions (ID_50_ values). Dashed horizontal lines correspond to the background values representing the limit of detection for neutralization assays. Significant differences between cohorts linked by horizontal lines are indicated by asterisks: p<0.05 = *, p<0.01 = **, p<0.001 = ***.

### mRNA-encoded S-EABR construct induces cell surface expression and eVLP budding

A key advantage of the EABR eVLP technology over existing nanoparticle-based vaccine approaches is that S-EABR constructs can be easily delivered as mRNA vaccines since both eVLP assembly and cell surface expression only require expression of a single genetically encoded component. While conventional COVID-19 mRNA vaccines induce antibody responses through cell surface expression of S protein (Figure 3A, top), mRNA-mediated delivery of an S-EABR construct could enhance B cell activation because S-EABR proteins will not only be expressed at the cell surface – they will also induce assembly of eVLPs that bud from the cell and distribute inside the body to activate immune cells (Figure 3A, bottom).

**Figure 3.**
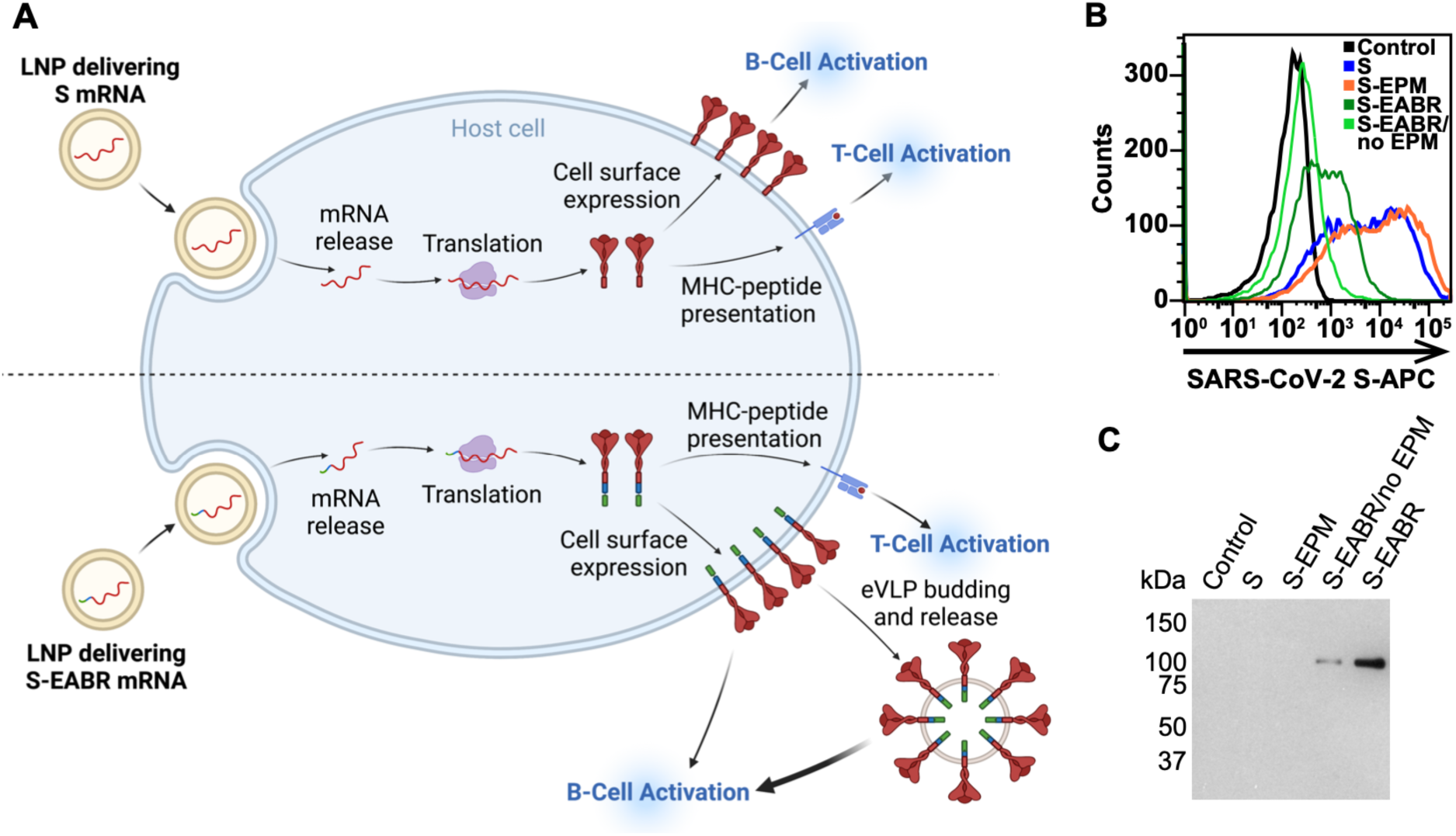
mRNA-mediated delivery of the S-EABR construct results in cell surface expression and eVLP assembly. (A) Schematic comparison of mRNA-LNP delivery of S (as in COVID-19 mRNA vaccines) (top) versus delivery of an S-EABR construct (bottom). Both approaches generate S peptides displayed on class I MHC molecules for CD8^+^ T cell recognition and result in presentation of S antigens on cell surfaces. The S-EABR approach also results in budding and release of eVLPs displaying S antigens. (B) Flow cytometry analysis of SARS-CoV-2 S cell surface expression on HEK293T cells that were untransfected (black) or transfected with mRNAs encoding S (blue), S-EPM (orange), S-EABR (dark green), or S-EABR/no EPM (light green) constructs. (C) Western blot analysis of eVLPs purified by ultracentrifugation on a 20% sucrose cushion from supernatants from the transfected cells in panel B. Purified eVLP samples were diluted 1:10.

To investigate whether genetic encoding of S-EABR eVLPs enhances the potency of a SARS-CoV-2 S-based mRNA vaccine, we started by synthesizing nucleoside-modified mRNAs encoding S, S-EABR, S-EPM, or S-EABR/no EPM. Cell surface expression and eVLP assembly were evaluated by flow cytometry and Western blot analysis 48 hours after in vitro transfection of mRNAs in HEK293T cells, demonstrating higher surface expression for S compared to the S-EABR fusion protein (Figure 3B). While addition of the EPM had little effect on S surface expression, removal of the EPM lowered surface levels for the S-EABR construct. Western blot analysis of supernatants confirmed that the S and S-EPM transfections did not generate detectable eVLPs in supernatants, whereas eVLPs were strongly detected in supernatants from S-EABR transfected cells (Figure 3C). eVLP production was decreased for S-EABR/no EPM, which together with the flow cytometry results (Figure 3B), suggests that EPM addition enhances both S-EABR cell surface expression and eVLP assembly.

The observed reduction in S cell surface expression in the S-EABR versus S mRNA transfections could be caused by lower overall cell surface expression of the S-EABR fusion protein, incorporation of S-EABR proteins into eVLPs that bud from the cell surface, or both. To evaluate these possibilities, we calculated approximate numbers of S trimers expressed from the S-EABR construct. Assuming that 3x10_6_ cells were transfected (6-well plate) and up to 1x10^5^ S trimers were expressed on the surface of each cell (based on the approximate number of B cell receptors on a B cell (Alberts et al., 2002)), transfected cell surfaces would contain ∼0.5 pmol or ∼70 ng of total S protein. Supernatant samples for Western blots were concentrated to a final volume of 200 µL of which 1.2 µL was loaded onto a gel. As the detection limit for S1 is ∼20 ng, the Western blot analysis suggested that purified S-EABR eVLPs from transfected cell supernatants contained at least ∼17 ng/µL S protein, corresponding to >3 µg S protein in the purified transfected cell supernatant. These calculations suggested that the observed reduction in cell surface expression for the S-EABR construct was at least partially caused by incorporation of S-EABR proteins into budding eVLPs that were released into the supernatant. Given that the estimated S protein content on released eVLPs exceeded the approximate amount of S protein presented on cell surfaces, it is possible that the S-EABR construct induces higher overall expression of S antigens compared to S for which expression is restricted to cell surfaces. Taken together, the mRNA transfection results demonstrate that the mRNA-encoded S-EABR construct enables dual presentation of S antigens on cell surfaces and released eVLPs.

### S-EABR mRNA-LNP elicit superior antibody titers compared to conventional vaccines

The effect of eVLP production on mRNA vaccine potency was evaluated in BALB/c mice by comparing mRNAs encoding S or S-EABR constructs that were encapsulated in LNP (Figure 4A). As described for preclinical studies of a COVID-19 mRNA vaccine in mice (Corbett et al., 2020), mRNA-LNP were administered intramuscularly (IM) at a dose of 2 µg mRNA on days 0 and 28. mRNA-LNP immunizations were also compared to purified S-EABR eVLPs that were injected IM in the presence of Addavax adjuvant. Antibody binding and neutralizing responses were evaluated by ELISAs and pseudovirus neutralization assays, respectively (Figures 4B-4H). After the prime, S and S-EABR mRNA-LNP elicited significantly higher antibody binding responses against the SARS-CoV-2 S protein than purified S-EABR eVLPs (Figure 4C). However, the highest neutralizing antibody titers were elicited by purified S-EABR eVLPs, which were significantly higher than titers elicited by the S mRNA-LNP (Figure 4D).

**Figure 4.**
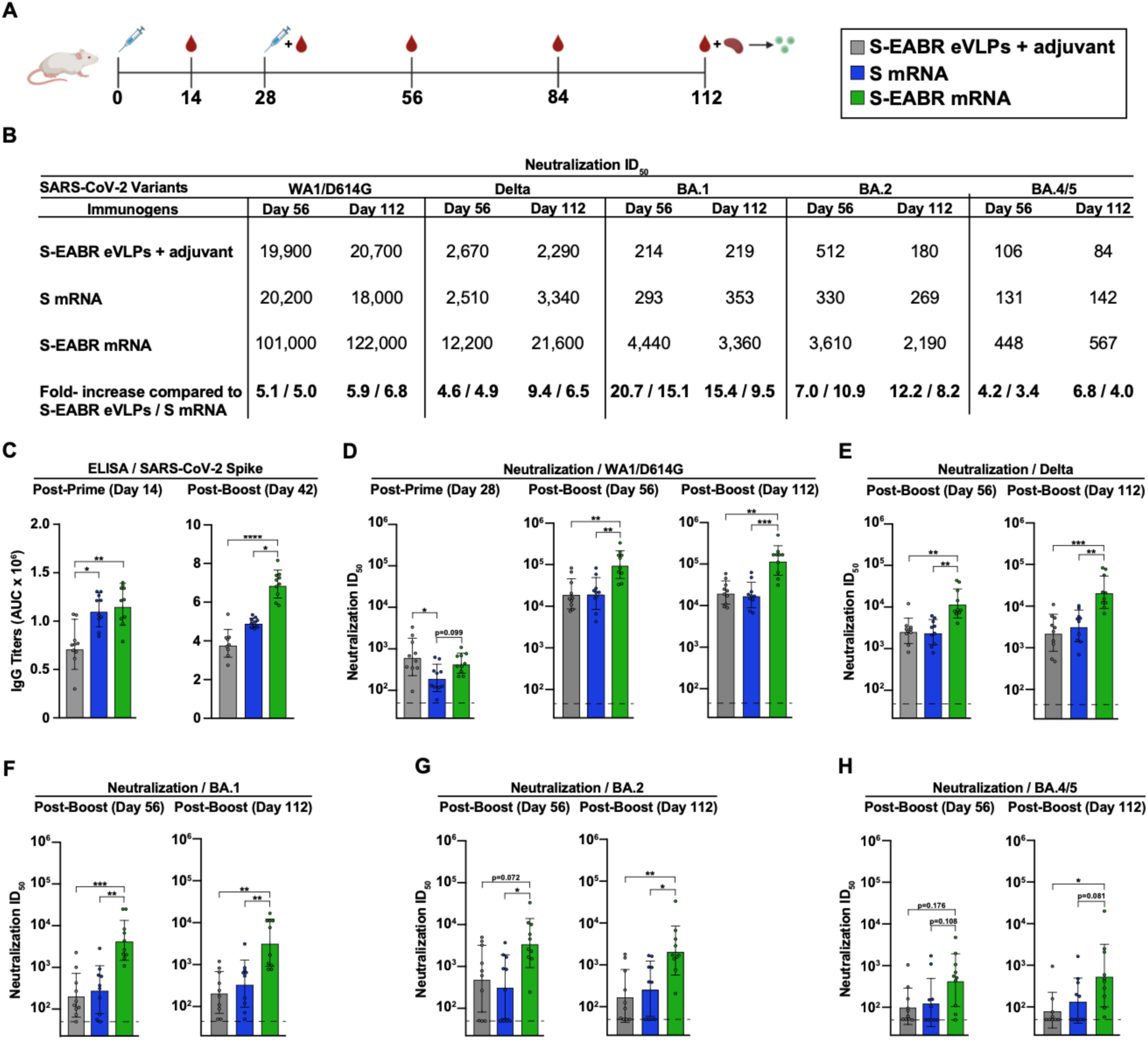
mRNA-LNP encoding S-EABR eVLPs induce potent antibody responses in mice. (A) Immunization schedule. BALB/c mice were immunized with purified S-EABR eVLPs (1 µg S protein) plus adjuvant (gray), 2 µg of mRNA-LNP encoding S (blue), or 2 µg of mRNA-LNP encoding S-EABR (green). On day 112, spleens were harvested from immunized mice for ELISpot analysis. (B) Neutralization data from indicated time points for antisera presented as geometric mean half-maximal inhibitory dilution (ID_50_) values against SARS-CoV-2 WA1/D614G, Delta, Omicron BA.1, Omicron BA.2, and Omicron BA.4/5 pseudoviruses. Bottom horizontal row shows the fold increases for geometric mean ID_50_ values for mice that received S-EABR mRNA-LNP compared to mice that received purified S-EABR eVLPs or S mRNA-LNP. (C) ELISA data from the indicated time points for antisera from individual mice (colored circles) presented as the geometric mean (bars) and standard deviation (horizontal lines). ELISAs evaluated binding of SARS-CoV-2 S trimers; results are shown as area under the curve (AUC). (D-H) Neutralization data from the indicated time points for antisera from individual mice (colored circles) presented as the geometric mean (bars) and standard deviation (horizontal lines). Neutralization results against SARS-CoV-2 WA1/D614G (D), Delta (E), Omicron BA.1 (F), Omicron BA.2 (G), and Omicron BA.4/5 (H) pseudoviruses are shown as ID_50_ values. Dashed horizontal lines correspond to the background values representing the limit of detection for neutralization assays. Significant differences between cohorts linked by horizontal lines are indicated by asterisks: p<0.05 = *, p<0.01 = **, p<0.001 = ***, p<0.0001 = ****.

After a boost immunization, S-EABR mRNA-LNP elicited significantly higher binding and neutralizing antibody titers than purified S-EABR eVLPs and S mRNA-LNP (Figures 4B-4D). Geometric mean neutralization titers measured for S-EABR mRNA-LNP were 5.1- and 5-fold higher than titers elicited by purified S-EABR eVLPs and S mRNA-LNP, respectively (Figures 4B and 4D). Three months post-boost (day 112), mean neutralization titers were 5.9- and 6.8-fold higher for S-EABR mRNA-LNP compared to purified S-EABR eVLPs and S mRNA-LNP, respectively, demonstrating that the increased serum neutralization activity was maintained (Figures 4B and 4D).

We also evaluated serum neutralizing activity against SARS-CoV-2 VOCs. S-EABR mRNA-LNP elicited 4.9- and 6.5-fold higher mean neutralizing responses against the Delta variant compared to S mRNA-LNP, as well as 4.6- and 9.4-fold higher titers compared to purified S-EABR eVLPs on days 56 and 112, respectively (Figures 4B and 4E). Against Omicron BA.1, neutralizing antibody responses dropped markedly for all groups, except for mice that received S-EABR mRNA-LNP, which elicited 15.1- and 9.5-fold higher neutralizing titers than S mRNA-LNP and 20.7- and 15.4-fold higher titers than purified S-EABR eVLPs on days 56 and 112, respectively (Figures 4B and 4F). Against Omicron BA.2, mean neutralization titers measured for mice that received S-EABR mRNA-LNP were also 10.9- and 8.2-fold higher compared to S mRNA-LNP and 7- and 12.2-fold higher compared to purified S-EABR eVLPs on days 56 and 112, respectively (Figures 4B and 4G). Compared to BA.2 titers, neutralizing antibody responses against the BA.4/5 variant decreased 4-8-fold for mice that received S-EABR mRNA-LNP, but titers were still 3.4- and 4-fold higher (but not statistically significant) compared to S mRNA-LNP and 4.2- and 6.8-fold higher compared to purified S-EABR eVLPs on days 56 and 112, respectively (Figures 4B and 4H). While neutralization titers of >1:400 against the BA.4/5 variant were measured for 7 of 10 mice that received S-EABR mRNA-LNP on day 56, such responses were only detected in 1 or 2 mice that received purified S-EABR eVLPs or S mRNA-LNP, respectively. Together, these results demonstrate that mRNA- mediated delivery of S-EABR eVLPs enhances the potency and breadth of humoral immune responses in mice compared to conventional mRNA and protein nanoparticle-based vaccine approaches. The observed improvements in neutralizing activity against Omicron-based VOCs were substantially larger than the 1.5-fold increases reported for recently approved bivalent mRNA booster shots (Khoury et al., 2022), suggesting that S-EABR mRNA-LNP-based booster immunizations could induce more effective and lasting immunity against Omicron-based and emerging VOCs than current COVID-19 vaccines.

### S-EABR mRNA-LNP induce potent T cell responses

On day 112 (3 months post-boost), splenocytes were isolated from immunized mice to analyze T cell responses by enzyme-linked immunosorbent spot (ELISpot) assays (Ranieri et al., 2014). Splenocytes were stimulated with a pool of SARS-CoV-2 S-specific peptides, and INF-γ and IL-4 secretion were measured to evaluate T cell activation. mRNA-LNP encoding S and S-EABR constructs induced potent INF-γ responses, consistent with the presence of S-specific cytotoxic CD8+ T cells and T helper 1 (T_H_1) cellular immune responses (Figure 5A). In contrast, INF-γ responses were almost undetectable for mice immunized with purified S-EABR eVLPs (Figure 5A). These results were expected as mRNA-LNP immunizations result in intracellular expression of S or S-EABR immunogens and MHC class I presentation of antigenic peptides that activate CD8+ T cells, which does not commonly occur for protein nanoparticle-based vaccines (Rock et al., 2016).

**Figure 5.**
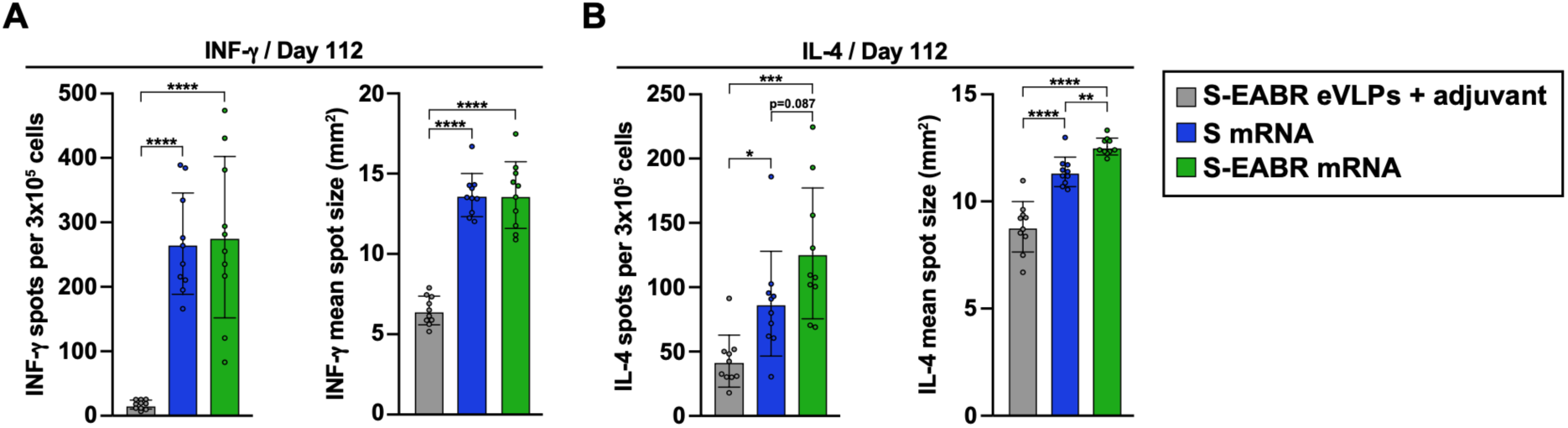
mRNA-LNP encoding S-EABR eVLPs induce potent T cell responses in mice. (A-B) ELISpot assay data for SARS-CoV-2 S-specific INF-γ (A) and IL-4 (B) responses of splenocytes from BALB/c mice that were immunized with purified S-EABR eVLPs (1 µg S protein) plus adjuvant (gray), 2 µg of mRNA-LNP encoding S (blue), or 2 µg of mRNA-LNP encoding S-EABR (green). Results are shown as spots per 3x10^5^ cells (left) and mean spot sizes (right) for individual mice (colored circles) presented as the mean (bars) and standard deviation (horizontal lines). Significant differences between cohorts linked by horizontal lines are indicated by asterisks: p<0.05 = *, p<0.01 = **, p<0.001 = ***, p<0.0001 = ****.

S-EABR mRNA-LNP induced significantly stronger IL-4 responses compared to S mRNA-LNP and purified S-EABR eVLPs (Figure 5B), consistent with potent T_H_2 cellular immune responses. While T_H_1- and T_H_2-biased responses were observed for S mRNA-LNP and purified S-EABR eVLPs, respectively, S-EABR mRNA-LNP induced a balanced T_H_1/T_H_2 response, thereby potently stimulating cellular and humoral immune responses. Thus, S-EABR mRNA-LNP retain the ability of conventional S mRNA-LNP to activate potent cytotoxic CD8+ T cell responses, while also potently activating T_H_2 CD4+ T cell responses to enhance humoral immune responses leading to increased antibody potency and breadth.

## Discussion

Here, we present a novel technology to generate eVLPs for vaccine and other applications. The approach harnesses the ESCRT pathway that is involved in cell division and viral budding (McCullough et al., 2018; Votteler and Sundquist, 2013) to drive assembly and release of eVLPs that present membrane proteins containing a cytoplasmic ESCRT-recruiting motif, the EABR sequence from the human centrosomal protein CEP55 (van der Horst et al., 2009). Our results demonstrate that the EABR-based platform produces eVLPs that incorporate higher levels of membrane antigens compared to approaches that require co-expression of the antigen with viral capsid proteins such as Gag or with the SARS-CoV-2 M, N, and E proteins. Purified S-EABR eVLPs elicited potent antibody responses against SARS-CoV-2 in mice that were similar in magnitude to those elicited by a 60-mer protein nanoparticle displaying S trimers. Compared to existing protein nanoparticle-based vaccine approaches, the EABR technology exhibits attractive manufacturing properties as (*i*) eVLP production requires expression of only a single component, (*ii*) transmembrane proteins are retained in their native membrane-associated conformation to ensure optimal protein expression and stability, and (*iii*) fully assembled eVLPs can be purified directly from culture supernatants without requiring detergent-mediated cell lysis and separation of membrane protein antigens from cell lysates. The lipid bilayer surrounding eVLPs also prevents off-target antibody responses against a nanoparticle scaffold that have been reported for protein nanoparticle-based immunogens (Kraft et al., 2022). Due to its modularity, flexibility, and versatility, the EABR technology could potentially be used to generate eVLPs presenting a wide range of surface proteins for vaccine and therapeutic applications.

To optimize the EABR technology, we evaluated several ESCRT-recruiting motifs for their ability to drive eVLP assembly, including viral late domains from EIAV, HIV-1, and EBOV. The EABR from CEP55 generated eVLPs 10-fold more efficiently than the EIAV late domain p9. The EABR binds to ESCRT proteins ALIX and TSG101 (Lee et al., 2008), while p9 binds only to ALIX (Fisher et al., 2007), suggesting that efficient eVLP assembly requires recruitment of both proteins. HIV-1 p6 contains motifs that interact with both TSG101 and ALIX (Fisher et al., 2007; Fujii et al., 2009), but S-p6 constructs did not induce detectable eVLP budding in our experiments, perhaps because reported affinities are relatively low (Fisher et al., 2007; Pornillos et al., 2002) compared to TSG101 and ALIX affinities reported for the EABR (Lee et al., 2008). eVLP production might be optimized by designing ESCRT-binding motifs with increased affinities for ESCRT proteins. We were able to enhance eVLP production by including an EPM derived from the FcgRII-B1 cytoplasmic tail (Miettinen et al., 1992) to reduce endocytosis of EABR-fusion proteins, which increased S-EABR cell surface expression and eVLP production.

An advantage of the EABR technology is that constructs can be easily delivered as mRNA vaccines since eVLP assembly requires expression of only a single component. This strategy results in presentation of viral surface antigens on the cell surface and on released eVLPs that could distribute throughout the body, thereby combining immune responses elicited by both conventional mRNA and protein nanoparticle-based vaccines. S-EABR mRNA-LNP elicited significantly higher binding and neutralizing antibody responses compared to conventional S-based mRNA-LNP analogous to current COVID-19 mRNA vaccines and to purified S-EABR eVLPs, suggesting that dual presentation of viral surface antigens on cell surfaces and eVLPs potentiates B cell activation. Presentation of viral surface antigens on cell surfaces alone potentially restricts expression for conventional mRNA vaccines due to a finite, and presumably limited, environment for insertion of both delivered and endogenous membrane proteins. Thus, combining cell surface expression and eVLP release for the S-EABR mRNA-LNP may increase overall presentation of viral surface antigens to the immune system. It is also possible that mRNA-mediated S-EABR eVLP production expands the biodistribution of viral surface antigens to more effectively engage B cells in lymph nodes distant from the injection site. The enhanced humoral immune responses elicited by S-EABR mRNA-LNP were consistent with potent T_H_2 cellular responses observed in S-EABR mRNA-LNP-immunized mice, which were more pronounced than in mice immunized with S mRNA-LNP or purified S-EABR eVLPs. Importantly, cytotoxic CD8+ T cell responses were maintained in S-EABR mRNA-LNP-compared to S mRNA-LNP-immunized animals. Thus, S-EABR mRNA-LNP potently stimulate both cellular and humoral immune responses.

The higher peak antibody levels elicited by the S-EABR mRNA-LNP would likely impact the durability of protective antibody responses. Notably, differences in serum antibody titers across different immunizations were maintained until three months post-boost, suggesting that antibody levels might contract at similar rates for the tested vaccine types. Hence, the elevated peak antibody titers elicited by the S-EABR mRNA-LNP could result in markedly prolonged periods of immune protection compared to conventional vaccine approaches, which could minimize the need for frequent booster immunizations. Long-term studies that monitor antibody levels for several months are needed to elucidate the relationship between peak antibody titers and durability of responses.

Two immunizations with S-EABR mRNA-LNP also elicited potent neutralizing antibody responses against SARS-CoV-2 Delta and Omicron-based VOCs, suggesting that higher antibody responses could lead to enhanced protection against viral escape variants. The conventional S-based mRNA-LNP immunization only elicited weak responses against Omicron-based VOCs, consistent with outcomes reported in humans in which weak Omicron-specific responses to WA1-based vaccines were enhanced after a 3^rd^ immunization (Barouch, 2022; Gruell et al., 2022). S-EABR mRNA-LNP elicited >10-fold higher neutralizing antibody titers against Omicron BA.1 and BA.2 VOCs compared to S mRNA-LNP after only two immunizations, suggesting that the simple addition of a short EABR-encoding sequence to the spike gene in current mRNA vaccines could have limited the global spread of Omicron-based VOCs. Our results also suggest that S-EABR mRNA-LNP-based booster immunizations would induce superior immunity against Omicron-based and emerging VOCs compared with current boosting strategies, as bivalent booster shots that contain ancestral and Omicron-based variants improve neutralizing antibody titers by only 1.5-fold compared to conventional booster shots (Khoury et al., 2022). Future studies need to investigate whether the observed increase in neutralization activity against Omicron-based VOCs results from higher overall antibody levels and/or increased antibody targeting of sub-immunodominant conserved epitopes on S trimer.

Enhanced antibody responses compared to S mRNA-LNP have also been reported for co-delivery of mRNAs encoding SARS-CoV-2 S, M, and E proteins, which should result in dual presentation of S on cell surfaces and released eVLPs (Lu et al., 2020). However, higher mRNA doses (10 µg) were needed to deliver all three mRNAs, and only modest improvements (∼2.5-fold) in neutralizing antibody titers were achieved. Our results showed that S-EABR eVLPs assemble more efficiently in vitro than eVLPs driven by co-expression of S, M, N, and E proteins, potentially explaining why S-EABR mRNA-LNP induced larger increases in antibody titers at lower doses. Co-delivery of multiple mRNAs also poses an obstacle for vaccine manufacturing, whereas COVID-19 and other mRNA vaccines could be easily modified to generate eVLPs by adding a short sequence containing EABR and EPM motifs to the cytoplasmic domains of the encoded immunogens. mRNA delivery of a trimerized RBD-ferritin fusion construct, which should result in secretion of non-enveloped ferritin nanoparticles displaying trimeric RBDs without cell surface expression of RBDs, has also been reported (Sun et al., 2021). This approach was not compared to a conventional S mRNA-LNP-based immunogen, highlighting the need for comparison studies of different vaccine approaches to elucidate the individual effects of antigen presentation on cell surfaces and virus-like nanoparticles on the magnitude and quality of immune responses.

In summary, we present a novel technology to efficiently generate eVLPs for vaccine and other therapeutic applications. We demonstrate that an mRNA vaccine encoding SARS-CoV-2 spike-EABR eVLPs elicits antibody responses with enhanced potency and breadth compared to conventional vaccine strategies in mice, which warrants further investigation in other preclinical animal models and humans as a vaccine strategy.

## Acknowledgements

We thank J. Vielmetter and the Caltech Protein Expression Center for assistance with protein production, K. Dam for biotinylated proteins for ELISAs, M. Anaya for a BirA expression plasmid, and J. Bloom (Fred Hutchinson) and P. Bieniasz (Rockefeller University) for neutralization assay reagents. We thank J. Keeffe, A. West, Y. Tam (Acuitas Therapeutics), C. Barnes (Stanford), H. Kleanthous (Bill and Melinda Gates Foundation), B. Wold, G. Tolomiczenko, and the Caltech Merkin Institute for Translational Research for helpful discussions and A. West for careful reading of the manuscript. Electron tomography was performed in the Caltech Cryo-EM Center with assistance from S. Chen. We thank Labcorp Drug Development–Antibody Reagents and Vaccines (Denver, PA) (formerly Covance, Inc.) for mouse immunization studies, BIOQUAL, Inc. (Rockville, MD) for PRNT50 assays, and R. Sukhovershin and RNAcore (Houston Methodist Research Institute) for synthesis of mRNAs and helpful discussion. Figures 1A and 3A were created with Biorender.com. This work was supported by the Bill and Melinda Gates Foundation INV-034638 (P.J.B.), the Caltech Merkin Institute (P.J.B.), the George Mason University Fast Grants (P.J.B.), the Rothenberg Innovation Initiative (RI^2^) (P.J.B.), and Wellcome Leap (P.J.B.). M.A.G.H. was supported by an NIH Director’s Early Independence Award (FAIN# DP5OD033362).

## Author contributions

M.A.G.H. and P.J.B. conceived the study, acquired funding, analyzed the data, and wrote the manuscript with contributions from other authors (Z.Y., P.J.C.L.). M.A.G.H. and K.E.H.T. generated, expressed, and evaluated EABR constructs by Western blot and flow cytometry analysis. M.A.G.H., K.E.H.T., P.N.P.G., L.M.K., and K.N.S. evaluated serum antibody responses from immunized mice by ELISA and neutralization assays. Z.Y. performed cryo-electron tomography and interpreted results. A.A.C. prepared S-mi3 immunogens for immunization studies in mice. W.J.M. and P.J.C.L. prepared mRNA-LNP for immunization studies in mice.

## Data availability

All data are available in the main text or the supplementary information. Materials are available upon request to the corresponding authors with a signed material transfer agreement. This work is licensed under a Creative Commons Attribution 4.0 International (CC BY 4.0) license, which permits unrestricted use, distribution, and reproduction in any medium, provided the original work is properly cited. To view a copy of this license, visit https://creativecommons.org/licenses/by/4.0/. This license does not apply to figures/photos/artwork or other content included in the article that is credited to a third party; obtain authorization from the rights holder before using such material.

## Supplemental Information

## Methods

### Design of EABR constructs

The EABR domain (residues 160-217) of the human CEP55 protein was fused to the C-terminus of the SARS-CoV-2 S protein (WA1/D614G) separated by a 4-residue (Gly)_3_Ser (GS) linker to generate S-EABR/no EPM. This construct contained the native furin cleavage site, 2P stabilizing mutations (Pallesen et al., 2017), and the C-terminal 21 residues were truncated to remove an ER-retention signal (McBride et al., 2007). The S-EABR construct was generated by inserting residues 243-290 of mouse FcgRII-B1 upstream of the 4-residue GS linker and the EABR domain. The S-EABR_min1_ and S-EABR_min2_ constructs encoded residues 170-217 and 170-208 of CEP55, respectively. EABR constructs were also generated for HIV-1 Env_YU2_ and human CCR5. S-p6, S-VP40_1-44_, and S-p9 were generated by replacing the EABR domain gene with sequences encoding HIV-1 p6 (isolate HXB2), EBOV VP40 (residues 1-44; Zaire EBOV), and EIAV p9 (strain Wyoming), respectively. The S-ferritin construct was designed as described (Powell et al., 2021) by fusing genes encoding the ectodomain of SARS-CoV-2 S WA1/D614G containing a furin cleavage site and 2P mutations, and *Helicobacter pylori* ferritin, separated by a 3-residue Ser-Gly-Gly linker. All constructs were cloned into the p3bNC expression plasmid.

### Production of EABR eVLPs

EABR eVLPs were generated by transfecting Expi293F cells (Gibco) cultured in Expi293F expression media (Gibco) on an orbital shaker at 37°C and 8% CO_2_. Gag-based eVLPs were produced by co-transfecting Expi293F cells with a plasmid expressing Rev-independent HIV-1 Gag-Pol (pHDM-Hgpm2 plasmid; PlasmID Repository, Harvard Medical School) and SARS-CoV-2 S, HIV-1 Env_YU2_, or CCR5, respectively, at a ratio of 1:1. SARS-CoV-2 M/N/E-based eVLPs were produced by co-transfecting Expi293F cells with plasmids expressing the SARS-CoV-2 M, N, E, and S proteins at a ratio of 1:1:1:1. To enable interactions between M, N, E, and S, we transfected full-length S with an untruncated cytoplasmic domain. 72 hours post-transfection, cells were centrifuged at 400 x g for 10 min, supernatants were passed through a 0.45 μm syringe filter and concentrated using Amicon Ultra-15 centrifugal filters with 100 kDa molecular weight cut-off (Millipore). eVLPs were purified by ultracentrifugation at 50,000 rpm (135,000 x g) for 2 hours at 4°C using a TLA100.3 rotor and a Optima^TM^ TLX ultracentrifuge (Beckman Coulter) on a 20% w/v sucrose cushion. Supernatants were removed and pellets were re-suspended in 200 μL sterile PBS at 4°C overnight. To remove residual cell debris, samples were centrifuged at 10,000 x g for 10 min and supernatants were collected. For in vivo studies and cryo-ET, eVLPs were further purified by SEC using a Superose 6 10/300 column (GE Healthcare) equilibrated with PBS. Peak fractions corresponding to S-EABR eVLPs were combined and concentrated to 250-500 μL in Amicon Ultra-4 centrifugal filters with 100 kDa molecular weight cut-off. Samples were aliquoted and stored at -20°C.

### Protein expression

Soluble SARS-CoV-2 S-6P trimers (WA1/D614G) (Hsieh et al., 2020) and RBDs were expressed as described (Cohen et al., 2022; Wang et al., 2022). Briefly, Avi/His-tagged proteins were purified from transiently-transfected Expi293F cells (Gibco) by nickel affinity chromatography and SEC (Barnes et al., 2020; Cohen et al., 2022; Wang et al., 2022). Peak fractions corresponding to S-6P or RBD proteins were pooled, concentrated, and stored at 4°C. Biotinylated proteins for ELISAs were generated by co-transfection of Avi/His-tagged S-6P and RBD constructs with a plasmid encoding an endoplasmic reticulum-directed BirA enzyme (kind gift from Michael Anaya, Caltech). S-6P constructs with a C-terminal SpyTag003 tag (Keeble et al., 2019) were expressed for covalent coupling to a 60-mer protein nanoparticle (SpyCatcher003-mi3) using the SpyCatcher-SpyTag system (Brune et al., 2016; Zakeri et al., 2012).

### Preparation of SpyCatcher003-mi3 nanoparticles

SpyCatcher003-mi3 (Cohen et al., 2021) displaying SpyTagged SARS-CoV-2 S-6P trimers were prepared as described (Cohen et al., 2021; Cohen et al., 2022). Briefly, SpyCatcher003-mi3 subunits with N-terminal 6xHis tags were expressed in BL21 (DE3)-RIPL *E. coli* (Agilent). Bacterial cell pellets were lysed using a cell disruptor in the presence of 2.0 mM PMSF (Sigma). Lysates were centrifuged at 21,000 x g for 30 min, and supernatants were collected and filtered through a 0.2 µm filter. SpyCatcher003-mi3 was purified by Ni-NTA chromatography using a pre-packed HisTrap^TM^ HP column (GE Healthcare), concentrated in Amicon Ultra-15 centrifugal filters with 30 kDa molecular weight cut-off (Millipore), and purified by SEC on a HiLoad 16/600 Superdex 200 column (GE Healthcare) equilibrated with TBS. S-mi3 nanoparticles were generated by incubating purified SpyCatcher003-mi3 with a 3-fold molar excess of purified SpyTagged S-6P trimer overnight at 4°C in TBS. Conjugated S-mi3 nanoparticles were separated from uncoupled S-6P trimers by SEC using a Superose 6 10/300 column (GE Healthcare) equilibrated with PBS. Fractions corresponding to conjugated S-mi3 were identified by sodium dodecyl sulfate polyacrylamide gel electrophoresis (SDS-PAGE) and pooled.

### Western blot analysis

The presence of SARS-CoV-2 S, HIV-1 Env_YU2_, and CCR5 on purified eVLPs was detected by Western blot analysis. Samples were diluted in SDS-PAGE loading buffer under reducing conditions, separated by SDS-PAGE, and transferred to nitrocellulose membranes (0.2 μm) (GE Healthcare). The following antibodies were used for detecting SARS-CoV-2 S, HIV-1 Env_YU2_, and CCR5: rabbit anti-SARS-CoV-2 S1 protein (PA5-81795; ThermoFisher) at 1:2,500, the human anti-HIV-1 Env broadly neutralizing antibody 10-1074 (Mouquet et al., 2012) (expressed in-house) at 1:10,000, rat anti-CCR5 (ab111300; Abcam) at 1:2,000, HRP-conjugated mouse anti-rabbit IgG (211-032-171; Jackson ImmunoResearch) at 1:10,000, HRP-conjugated goat anti-human IgG (2014-05; Southern Biotech) at 1:8,000, and HRP-conjugated mouse anti-rat IgG (3065-05; Southern Biotech) at 1:10,000. Protein bands were visualized using ECL Prime Western Blotting Detection Reagent (GE Healthcare).

For in vivo studies, the amount of SARS-CoV-2 S on S-EABR eVLPs was determined by quantitative Western blot analysis. Various dilutions of SEC-purified S-EABR eVLP samples and known amounts of soluble SARS-CoV-2 S1 protein (Sino Biological) were separated by SDS-PAGE and transferred to nitrocellulose membranes (GE Healthcare). SARS-CoV-2 S was detected as described above. Band intensities of the SARS-CoV-2 S1 standards and S-EABR eVLP sample dilutions were measured using ImageJ to determine S concentrations. The S1 protein concentrations determined for S-EABR samples were multiplied by a factor of 1.8 to account for the difference in molecular weight between S1 and the full-length S protein.

### Cryo-ET of S-EABR eVLPs

SEC-purified S-EABR eVLPs were prepared on grids for cryo-ET using a Mark IV Vitrobot (ThermoFisher Scientific) operated at 21°C and 100% humidity. 2.5 μL of sample was mixed with 0.4 μL of 10 nm fiducial gold beads (Sigma-Aldrich) and applied to 300-mesh Quantifoil R2/2 grids, blotted for 3.5 s, and then plunge-frozen in liquid ethane cooled by liquid nitrogen. Image collections were performed on a 300 kV Titan Krios transmission electron microscope (ThermoFisher Scientific) operating at a nominal 42,000x magnification. Tilt series were collected on a K3 direct electron detector (Gatan) with a pixel size of 2.15 Å•pixel^-1^ using SerialEM software (Mastronarde, 2005). The defocus range was set to -5 to -8 μm and a total of 120 e^-^ • Å^-2^ per tilt series. Images were collected using a dose-symmetric scheme (Hagen et al., 2017) ranging from -60° to 60° with 3° intervals. Tomograms were aligned and reconstructed using IMOD (Mastronarde and Held, 2017).

To build a model of an S-EABR eVLP, coordinates of a SARS-CoV-2 S trimer (PDB 6VXX) were fit into spike densities in the reconstructed tomograms using ChimeraX (Goddard et al., 2018). Positions and orientations of the S protein were adjusted in a hemisphere of the eVLP in which the spike density was of higher quality. A 55 nm sphere was adapted from a cellPACK model (cellPACK ID: HIV-1_0.1.6_6) (Johnson et al., 2015; Johnson et al., 2014) and added to the model to represent the eVLP membrane surface.

### Neutralization assays

Lentivirus-based SARS-CoV-2 pseudoviruses were generated as described (Crawford et al., 2020; Robbiani et al., 2020) using S proteins from the WA1/D614G, Delta, Omicron BA.1, Omicron BA.2, and Omicron BA.4/5 variants in which the C-terminal 21 residues of the S protein cytoplasmic tails were removed (Crawford et al., 2020). Serum samples from immunized mice were heat-inactivated for 30 min at 56°C. Three-fold serial dilutions of heat-inactivated samples were incubated with pseudoviruses for 1 hour at 37°C, followed by addition of the serum-virus mixtures to pre-seeded HEK293T-ACE2 target cells. After 48-hour incubation at 37°C, BriteLite Plus substrate (Perkin Elmer) was added and luminescence was measured. Half-maximal inhibitory dilutions (ID_50_s) were calculated using 4-parameter non-linear regression analysis in AntibodyDatabase (West et al., 2013) and ID_50_ values were rounded to three significant figures.

PRNT_50_ (50% plaque reduction neutralization test) assays with authentic SARS-CoV-2 virus were performed in a biosafety level 3 facility at BIOQUAL, Inc. (Rockville, MD) as described (Haun et al., 2020). Mouse sera from day 56 post-immunization were diluted 1:20 and then 3-fold serially diluted in culture media (DMEM + 10% FBS + Gentamicin). The diluted samples were incubated with 30 plaque-forming units of wild-type SARS-CoV-2 (USA-WA1/2020, BEI Resources NR-52281; Beta variant, Isolate hCoV-19/South Africa/KRISP-K005325/2020, BEI Resources NR-54009; Delta variant, isolate hCoV-19/USA/MD-HP05647/2021 BEI Resources NR-55674) for 1 hour at 37°C. Samples were then added to a confluent monolayer of Vero/TMPRSS2 cells in 24-well plates for 1 hour at 37°C in 5% CO_2_. 1 mL of culture media with 0.5% methylcellulose was added to each well and plates were incubated for 3 days at 37°C in 5% CO_2_. Plates were fixed with ice cold methanol at -20°C for 30 min. Methanol was discarded and plates were stained with 0.2% crystal violet for 30 min at room temperature. Plates were washed once with water and plaques in each well were counted. TCID_50_ values were calculated using the Reed-Muench formula (Reed and Muench, 1938).

### ELISAs

Pre-blocked streptavidin-coated Nunc® MaxiSorp™ 384-well plates (Sigma) were coated with 5 µg/mL biotinylated S-6P or RBD proteins in Tris-buffered saline with 0.1% Tween 20 (TBS-T) and 3% bovine serum albumin (BSA) for 1 hour at room temperature. Serum samples from immunized mice were diluted 1:100, 4-fold serially diluted in TBS-T/3% BSA, and then added to plates. After a 3-hour incubation at room temperature, plates were washed with TBS-T using an automated plate washer. HRP-conjugated goat anti-mouse IgG (715-035-150; Jackson ImmunoResearch) was diluted 1:100,000 in TBS-T/3% BSA and added to plates for 1 hour at room temperature. After washing with TBS-T, plates were developed using SuperSignal^TM^ ELISA Femto Maximal Signal Substrate (ThermoFisher) and absorbance was measured at 425 nm. Area under the curve (AUC) calculations for binding curves were performed using GraphPad Prism 9.3.1 assuming a one-site binding model with a Hill coefficient as described (Cohen et al., 2021).

### mRNA synthesis

Codon-optimized mRNAs encoding SARS-CoV-2 S, S-EPM, S-EABR/no EPM, and S-EABR constructs were synthesized by RNAcore (https://www.houstonmethodist.org/research-cores/rnacore/) using proprietary manufacturing protocols. mRNAs were generated by T7 RNA polymerase-mediated in vitro transcription reactions using DNA templates containing the immunogen open reading frame flanked by 5′ untranslated region (UTR) and 3′ UTR sequences and terminated by an encoded polyA tail. CleanCap 5’ cap structures (TriLink) were incorporated into the 5′ end co-transcriptionally. Uridine was completely replaced with N1-methyl-pseudouridine to reduce immunogenicity (Kariko et al., 2008). mRNAs were purified by oligo-dT affinity purification and high-performance liquid chromatography (HPLC) to remove double-stranded RNA contaminants (Kariko et al., 2011). Purified mRNAs were stored at –80 °C.

### mRNA transfections

For mRNA transfections, 10^6^ HEK293T cells were seeded in 6-well plates. After 24 hours, cells were transfected with 2 µg mRNA encoding SARS-CoV-2 S, S-EPM, S-EABR/no EPM, or S-EABR constructs using Lipofectamine^TM^ MessengerMax^TM^ transfection reagent (ThermoFisher). 48 hours post-transfection, supernatants were collected and purified for Western blot analysis. Cells were gently detached by pipetting and resuspended in 500 µL PBS. 100 µL were transferred into Eppendorf tubes for flow cytometry analysis of S cell surface expression. Cells were stained with the SARS-CoV-2 antibody C119 (Robbiani et al., 2020) at 5 µg/mL in PBS+ (PBS supplemented with 2% FBS) for 30 min at room temperature in the dark. After two washes in PBS+, samples were stained with an Alexa Fluor® 647-conjugated anti-human IgG secondary antibody (A21445; Life Technologies) at a 1:2,000 dilution in PBS+ for 30 min at room temperature in the dark. After two washes in PBS+, cells were resuspended in PBS+ and analyzed by flow cytometry (MACSQuant, Miltenyi Biotec). Results were plotted using FlowJo 10.5.3 software.

### LNP encapsulation of mRNAs

Purified N1-methyl-pseudouridine mRNA was formulated in LNP as previously described (Pardi et al., 2015). In brief, 1,2-distearoyl-*sn*-glycero-3-phosphocholine, cholesterol, a PEG lipid, and an ionizable cationic lipid dissolved in ethanol were rapidly mixed with an aqueous acidic solution containing mRNA using an in-line mixer. The ionizable lipid and LNP composition are described in the international patent application WO2017075531(2017). The post in-line solution was dialyzed with PBS to remove the ethanol and displace the acidic solution. Subsequently, LNP was measured for size (60-65 nm) and polydispersity (PDI < 0.075) by dynamic light scattering (Malvern Nano ZS Zetasizer). Encapsulation efficiencies were >97% as measured by the Quant-iT Ribogreen Assay (Life Technologies).

### Immunizations

All animal procedures were performed in accordance with IACUC-approved protocols. 7-8 week-old female C57BL/6 or BALB/c mice (Charles River Laboratories) were used for immunization experiments with cohorts of 8-10 animals per group. 0.1 µg of protein-based immunogens, including soluble S trimer, S-mi3, and purified S-EABR eVLPs, were administered to C57BL/6 mice by subcutaneous (SC) injections on days 0 and 28 in the presence of Sigma adjuvant system (Sigma). 2 µg of S and S-EABR mRNA-LNP were administered to BALB/c mice by intramuscular (IM) injections on days 0 and 28. To compare mRNA- and protein-based immunogens, 1 µg purified S-EABR eVLPs were administered IM in the presence of 50% v/v AddaVax^TM^ adjuvant (Invivogen). Serum samples for ELISAs and neutralization assays were obtained on indicated days.

### ELISpot assays

Animals were euthanized on day 112 and spleens were collected. Spleens were homogenized using a gentleMACS Octo Dissociator (Miltenyi Biotec). Cells were passed through a 70 µm tissue screen, centrifuged at 1,500 rpm for 10 min, and resuspended in CTL-Test^TM^ media (ImmunoSpot) containing 1% GlutaMAX^TM^ (Gibco) for ELISpot analysis to evaluate T cell responses. A PepMix^TM^ pool of 315 peptides (15-mers with 11 amino acid overlap) derived from the SARS-CoV-2 S protein (JPT Peptide Technologies) was added to mouse IFN-g/IL-4 double-color ELISpot plates (ImmunoSpot) at a concentration of 2 µg/mL. 300,000 cells were added per well, and plates were incubated at 37°C for 24 hours. Biotinylated detection, streptavidin-alkaline phosphatase (AP), and substrate solutions were added according to the manufacturer’s guidelines. Plates were gently rinsed with water three times to stop the reactions. Plates were air-dried for two hours in a running laminar flow hood. The number of spots and the mean spot sizes were quantified using a CTL ImmunoSpot S6 Universal-V Analyzer (Immunospot).

### Statistical analysis

Titer differences between immunized groups of mice for ELISAs and neutralization assays were evaluated for statistical significance using the non-parametric Kruskal-Wallis test followed by Dunn’s multiple comparison post hoc test calculated using Graphpad Prism 9.3.1. For ELISpot results, statistically significant differences between immunized groups of mice were determined using analysis of variance (ANOVA) test followed by Tukey’s multiple comparison post hoc test calculated using Graphpad Prism 9.3.1.

---

**Figure S1.**
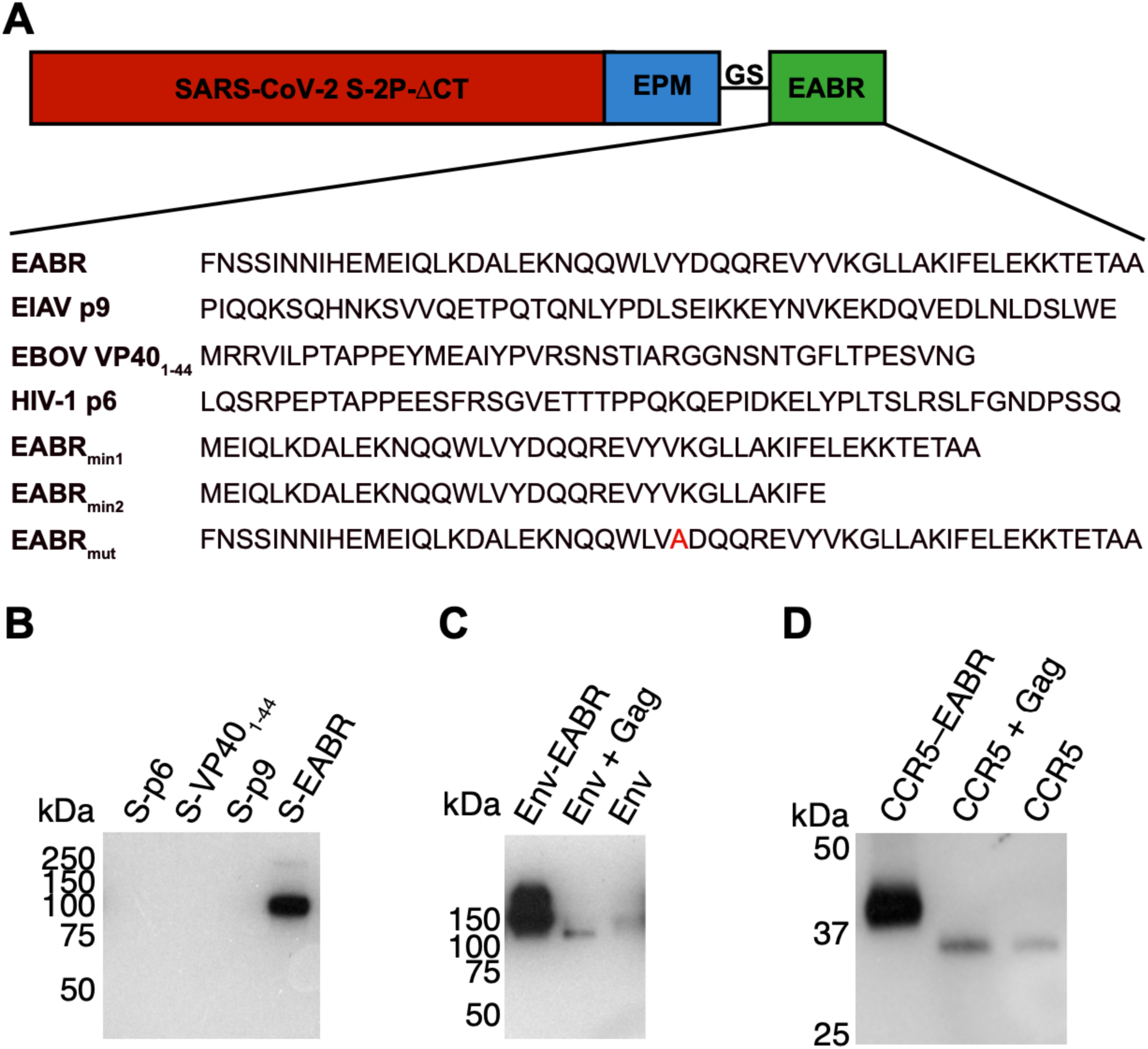
Comparison of EABR-related sequence insertions in the cytoplasmic tail of SARS-CoV-2 S, related to Figure 1. (A) Top: Schematic of different S-EABR constructs that were compared for their ability to induce eVLP assembly. EPM = Endocytosis prevention motif. GS = (Gly)_3_Ser linker. EABR = ESCRT- and ALIX-binding region. Bottom: Amino acid sequences of EABR portion of different constructs. (B) Western blot analysis of SARS-CoV-2 S1 protein levels on eVLPs purified by ultracentrifugation on a 20% sucrose cushion from transfected Expi293F cell culture supernatants. Cells were transfected with S-p6, S-VP40_1-44_, S-p9, or S-EABR constructs. Purified eVLP samples were diluted 1:400. (C) Western blot analysis comparing HIV-1 Env_YU2_ levels in eVLP samples purified from transfected Expi293F cell culture supernatants. Cells were transfected with plasmids encoding Env-EABR, Env plus HIV-1 Gag, or Env alone. Purified eVLP samples were diluted 1:200. (D) Western blot analysis comparing CCR5 levels in eVLP samples purified from transfected Expi293F cell culture supernatants. Cells were transfected with plasmids encoding CCR5-EABR, CCR5 plus HIV-1 Gag, or CCR5 alone. Purified eVLP samples were diluted 1:200. The migration difference between CCR5-EABR and CCR5 is due to addition of the EABR sequence (∼7 kDa) that increases its molecular mass.

**Figure S2.**
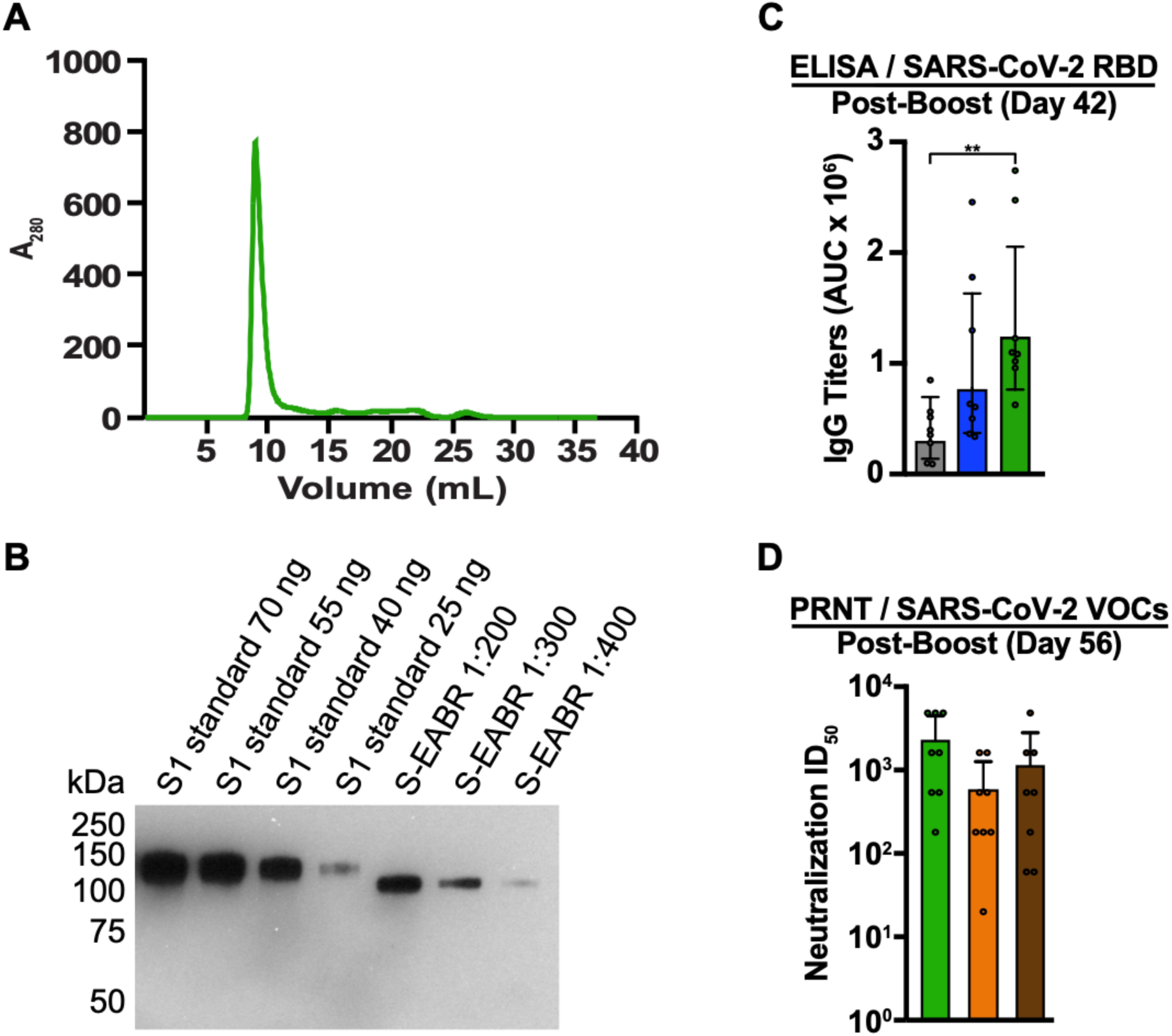
Purified S-EABR eVLPs induce potent antibody responses in mice, related to Figure 2. (A) Size exclusion chromatogram of S-EABR eVLPs purified by ultracentrifugation on a 20% sucrose cushion. (B) Quantitative Western blot comparing indicated amounts of SARS-CoV-2 S1 standards (lanes 1-4) and various dilutions of purified S-EABR eVLPs (lanes 5-7) to determine S protein concentrations in eVLP samples. The S1 standard protein (Sino Biological) was biotinylated and contained a polyhistidine tag, which resulted in a difference in apparent molecular weights for the S1 standards and the S-EABR construct. Band intensities of S1 standards and S-EABR eVLP sample dilutions were measured using ImageJ to determine S concentrations. (C) ELISA data from day 42 for antisera from individual mice (colored circles) immunized with soluble S (purified S trimer) (gray), S-mi3 (S trimer ectodomains covalently attached to mi3, a 60-mer protein nanoparticle) (blue), or S-EABR eVLPs (green). Results are shown as area under the curve (AUC) and presented as the geometric mean (bars) and standard deviation (horizontal lines). Significant differences between cohorts linked by horizontal lines are indicated by asterisks: p<0.05 = *, p<0.01 = **. (D) PRNT assay results from day 56 for antisera from individual mice (colored circles) immunized with S-EABR eVLPs. Results against the SARS-CoV-2 WA1 (green), Beta (orange), and Delta (brown) variants are shown as TCID_50_ values (Reed and Muench, 1938) and presented as the geometric mean (bars) and standard deviation (horizontal lines).

